# Resting-State Brain Fluctuation and Functional Connectivity Dissociate Moral Injury from Posttraumatic Stress Disorder

**DOI:** 10.1101/442327

**Authors:** Delin Sun, Rachel D. Phillips, Hannah L. Mulready, Stephen T. Zablonski, Jessica A. Turner, Matthew D. Turner, Kathryn McClymond, Jason A. Nieuwsma, Rajendra A. Morey

**Author notes:** Corresponding author: Delin Sun, Ph.D. Duke-UNC Brain Imaging and Analysis Center, 40 Duke Medicine Circle, Room 414, Durham, NC 2 7 710 USA, Facsimile: 1-919-416-5912.

## Abstract

Moral injury is closely associated with posttraumatic stress disorder (PTSD) and is characterized by disturbances in social and moral cognition. Little is known about the neural underpinnings of moral injury, and whether the neural correlates are different between moral injury and PTSD.

A sample of 26 US military veterans (2 females; 28~55 years old) were investigated to determine how moral injury experiences and PTSD symptoms are differentially related to spontaneous fluctuations indexed by low frequency fluctuation (ALFF) as well as functional connectivity during resting-state functional magnetic resonance imaging (fMRI) scanning.

ALFF in the left inferior parietal lobule (L IPL) was positively associated with moral injury sub-scores of transgressions, negatively associated with sub-scores of betrayals, and not related with PTSD symptoms. Moreover, functional connectivity between the L IPL and bilateral precuneus was positively related with PTSD symptoms and negatively related with moral injury total scores.

Our results provide the first evidence that moral injury and PTSD have dissociable neural underpinnings, and behaviorally distinct sub-components of moral injury are different in neural responses. The findings increase our knowledge of the neural distinctions between moral injury and PTSD and may contribute to developing nosology and interventions for military veterans afflicted with moral injury.

## Introduction

Morally judicious behavior is essential to the fabric of human societies and social relationships. Moral injury refers to disturbances experienced by combat veterans related to guilt, shame, and betrayal arising from violations of their moral code [1–3]. It may arise from specific acts, such as killing in combat (e.g., killing innocent civilians), but may also be generated by a broader experience that violates deeply held moral and ethical beliefs and expectations. Individual soldiers are left to make sense of their own actions and the actions of others, to integrate those actions with their existing moral and ethical frameworks, and to manage emotional responses prompted by the relative congruence or incongruence between past moral beliefs and recent actions. The inability to integrate long-held ethical worldviews with specific personal actions may lead to ongoing psychological distress manifested by specific behavioral problems [1; 4].

Both moral injury and military-related PTSD are associated with consequences of participation in warfare. A required criterion for a PTSD diagnosis is participating in a traumatic event; prevailing models of PTSD are predicated on exposure to life-threatening events and are predominantly studied as disorders of fear processing [5; 6]. Profound moral injury, on the other hand, may be experienced without directly participating in a traumatic event, and models of moral injury are related to disturbances in social and moral cognition [2]. Our goal was to investigate the hotly debated comparisons between moral injury and PTSD by probing the relevant neural systems [1; 7].

Psychometric evaluation of moral injury by the self-report Moral Injury Events Scale (MIES) developed by Nash et al. [8], shows that contributory events are captured by two latent factors: perceived transgressions by self or others (the transgression subscale), and perceived betrayals by others (betrayal subscale). The transgression subscale includes witnessing acts of commission, distress resulting from others′ acts of commission, and perpetration of or distress due to acts of commission/omission. An example would be a soldier who kills an unarmed civilian who was mistakenly perceived to be armed. On the other hand, the betrayal subscale measures perceived betrayals by previously trusted military leaders, fellow service members, and nonmilitary others (e.g., a spouse). For instance, a patriotic soldier in a battle may begin to wonder whether the war is not as justified as the leaders have declared.

Both moral transgression [3] and betrayal [9] have been linked to feelings of guilt and shame, which are associated with the ability to understand the social consequences of one′s own behavior as judged by others. In this sense, they are related with both self-referential and theory of mind (ToM) processes. Self-referential processing refers to functions for decoding information about oneself [10], while ToM refers to the ability to assign and attribute mental states to both self and others [11].

Both self-referential and ToM processes are associated with functions in default mode network (DMN) [12], which involves brain areas including medial prefrontal cortex (mPFC), posterior cingulate cortex (PCC), and inferior parietal lobule (IPL) [13]. The DMN is preferentially active when individuals are daydreaming, mind-wandering, engaged in internally focused tasks including retrieving autobiographical memory, envisioning the future, or thinking about others [14]. The neural correlates of moral processing largely overlap with the DMN [15].

Our previous work studying brain responses to guilt scenarios showed that the guilt ratings were positively associated with activations in dorsal mPFC and supramarginal gyrus that is included in IPL [16]. Roth et al. [17] investigated the neural correlates of autographical recall about shame and found that shame versus neutral condition elicited stronger activation in mPFC and PCC as well as weaker activation in IPL. Interestingly, studies on shame and guilt [15] have also reported findings in amygdala and dorsal anterior cingulate cortex (dACC) that are hyper-responsive in PTSD [18]. For instance, Pulcu et al. [19] detected increased amygdala response to shame in remitted major depressive disorder. For another example, Wagner et al. [20] found that guilt in healthy subjects elicited stronger activations in dACC and amygdala than shame. These previous studies explored the neural correlates of shame and guilt, which are the core components of moral injury [2; 3].

In the present study, we investigated the relationship between clinical measures of moral injury/PTSD and brain responses including both spontaneous fluctuation and functional connectivity (FC) during resting-state functional magnetic resonance imaging (rs-fMRI) scanning. The rs-fMRI does not measure responses to explicit tasks and is thus convenient for investigating the brain’s functional organization in patients with psychiatric and behavioral disorders. A number of studies have demonstrated that rs-fMRI data predict following behavioral performance in explicit tasks [21–23]. Here we measured the intensity of spontaneous fluctuations in the brain using two methods: the amplitude of low frequency fluctuations (ALFF) [24] and fractional ALFF (fALFF) [25]. These are common analysis approaches for spontaneous neural activity during rs-fMRI and have been widely employed to investigate the neural underpinnings of various mental psychiatric and behavioral disorders [25; 26]. ALFF and fALFF are positively correlated with other measures of spontaneous fluctuations such as regional homogeneity [27] and regional connectivity [28], thus we limited analyses to ALFF and fALFF methods for simplicity. Moreover, we measured functional connectivity based on the correlation between the blood-oxygen-level dependent (BOLD) time course of the seed region and that of all other areas in the brain [29]. The functional connectivity method has also been widely used in studies on psychiatric disorders [30].

Studies on shame and guilt using rs-fMRI techniques are scarce, while previous work showed that PTSD patients compared to trauma-exposed and non-trauma controls are associated with altered spontaneous brain activity during rs-fMRI in several areas including those within DMN such as mPFC, PCC [31], and IPL [32]. Moreover, altered resting-state functional connectivity of amygdala was reported in PTSD [33], and the functional connectivity patterns in DMN were also found to relate with PTSD symptoms [34; 35]. Based on previous task- or resting-state based fMRI studies of moral processing and PTSD, we anticipated to find neural correlates of moral injury or PTSD indexed by either ALFF/fALFF in regions of interest (ROIs) including the DMN areas as well as amygdala and dACC [32] or functional connectivity between these ROIs (seeds) and the rest of the brain. We also examine the relationships between resting-state brain responses and transgression- or betrayal-related sub-scores of the MIES.

## Methods and Materials

### Participants and Procedure

Detailed demographic and clinical information are described in **Table 1**. Participants were recruited from Iraq and Afghanistan era military service members in the VA Mid-Atlantic MIRECC Post-Deployment Mental Health Repository [36]. The present moral injury study combined data from 26 participants who participated in two post-repository studies of combat-exposed veterans focused on [1] moral injury and [2] rs-fMRI. In study [1] approximately 300 participants completed questionnaire packets by mail that included MIES to assess moral injury [8], depressive symptoms using the Beck Depression Inventory-II (BDI-II) [37], and combat exposure with the Combat Exposure Scale (CES) [38]. In study [2] participants completed a battery of measures, including determination of PTSD diagnosis using the Clinician Administered PTSD Scale (CAPS) [39; 40] based on symptoms experienced in the past month. Eleven out of 26 participants were diagnosed with PTSD. Descriptions of the CAPS, MIES, BDI-II and CES are in the supplementary materials. To be eligible for the present study, participants needed to have deployed to a combat zone and could not have a DSM-IV based diagnosis of psychosis. All participants in this study provided verbal informed consent to participate in procedures reviewed and approved by the Institutional Review Boards at Duke University and the Durham VA Medical Center.

**Table 1.** Demographic and clinical information (N=26, 2 females).

|  | Mean | STD | Range | Max Range |
| --- | --- | --- | --- | --- |
| Age (yrs) | 43.5 | 8.8 | 28-55 | NA |
| MIES-transgression | 14.2 | 7.2 | 6-30 | 6-36 |
| MIES-betrayal | 6.3 | 4.5 | 3-18 | 3-18 |
| MIES-total | 20.5 | 10.5 | 9-44 | 9-54 |
| CAPS | 28.5 | 32.9 | 0-100 | 0-136 |
| BDI-II | 12.5 | 14.4 | 0-54 | 0-69 |
| CES | 10.6 | 9.94 | 0-29 | 41 |

Note: MIES transgression and MIES betrayal were measured by the Moral Injury Events Scale (MIES) [8]. PTSD symptoms were measured through the Clinician Administered PTSD Scale (CAPS) [59]. Depressive symptoms were measured by the Beck Depression Inventory-II [37]. Combat exposure was measured by the Combat Exposure Scale (CES) [38]. STD, standard deviation; Range, range of values in our sample; Max Range, possible range according to the questionnaires and scales.

### Brain Image Acquisition, Preprocessing and ROI Selection

The detailed information of brain image acquisition, preprocessing and ROI (also seed) selection could be found in the supplementary document. We here employed 7 ROIs: mPFC, PCC, Left/Right IPL, Left/Right Amygdala, and dACC, as shown in **Fig. 1**.

**Figure 1.**
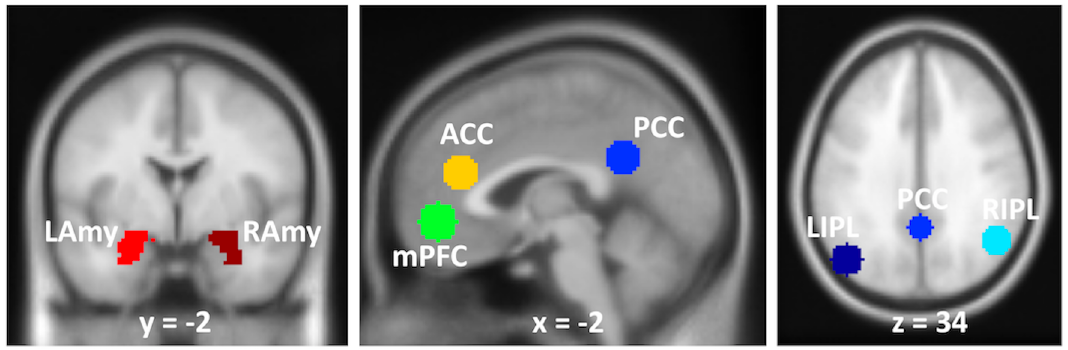
Regions of interest (ROIs) in the default mode network (DMN). mPFC, medial prefrontal cortex; PCC, posterior cingulate cortex; L/R IPL, left/right inferior parietal lobule; ACC, anterior cingulate cortex; L/R Amy, left/right amygdala.

### Statistical Models

Two statistical models were employed to fully understand the relationship between resting-state brain responses and self-reports of moral injury as well as PTSD. In both models, the variables including age, sex, BDI-II, CES scores, and study protocols were entered as covariates of no interest. In **Model I**, we investigated the neural correlates of either MIES-total (the sum of MIES-transgression and MIES-betrayal scores) or CAPS scores. MIES-total score was listed as a covariate of no interest when studying the neural correlates of CAPS, and CAPS score was a covariate of no interest when studying the neural correlates of MIES-total.

**Model I** may have overlooked the differences of moral injury subscales (transgression and betrayal). To further understand the neural underpinnings of moral injury subscales, we investigated in **Model II** the neural correlates of either MIES-transgression, MIES-betrayal or CAPS scores. Similar to Model I, when investigating the neural correlates of one of the clinical measures, the other measures served as the covariates of no interest.

To investigate whether the correlations were statistically different, we employed the Williams’s t-test [41], which is appropriate for the comparison between two non-independent correlations with a variable in common.

### ALFF and fALFF ROI Analyses

The mean ALFF and fALFF values from each of the seven ROIs were extracted using the MarsBar toolbox (http://marsbar.sourceforge.net). The MATLAB partial correlation function was utilized to control for the effects of covariates of no interest. The FDR method [42] was applied to correct for the number of correlations (total number = 28) across 7 ROIs, 2 clinical measures (MIES-total and CAPS) and 2 brain activity measures (ALFF and fALFF) in **Model I**. It was also employed for the number of correlations (total number = 42) across 7 ROIs, 3 clinical measures (MIES-transgression, MIES-betrayal and CAPS) and 2 brain activity measures (ALFF and fALFF) in **Model II**.

### ALFF and fALFF Whole-Brain Analyses

We employed two multiple regression models (corresponding to Model I and II, respectively) to investigate the relationship between ALFF/fALFF per voxel and each of the variables of interest after controlling the effects of all the other variables. Results were thresholded at *p* < 0.001 uncorrected and survived *p* < 0.05 cluster-extent size false discovery rate (FDR) correction.

### Seed-Based functional connectivity Whole-Brain Analyses

We employed two multiple regression models (corresponding to Model I and II, respectively) to investigate how the functional connectivity between a seed (one of 7 ROIs mentioned above) and voxels in the rest of the brain was related with each of the variables of interest after controlling the effects of all the other covariates. Results were thresholded at *p* < 0.001 uncorrected and survived *p* < 0.007 (< 0.05/7 given that there were 7 seeds) cluster-size false discovery rate (FDR) correction. If a clinical measure was found significantly correlate with functional connectivity between a seed and a target brain area, further voxel-wised analyses were conducted to test whether the other clinical measures also correlate with functional connectivity between the same pair of seed and target area. The findings were thresholded at *p* < 0.001 uncorrected and survived *p* < 0.05 small volume corrected (SVC) within the target area.

## Results

### ALFF and fALFF ROI Findings

For **Model I**, partial correlations between MIES-total or CAPS scores and average ALFF or fALFF in all of the ROIs are shown in **Table 2**. No results from **Model I** survived FDR corrections.

**Table 2.** Partial correlations between MIES-total/PTSD and average ALFF/fALFF in ROIs

| ROI | MIES-total |  |  | CAPS |  |  |
| --- | --- | --- | --- | --- | --- | --- |
|  | *R* | *p* | *adj_p* | *R* | *p* | *adj_p* |
| **ALFF** |  |  |  |  |  |  |
| mPFC | 0.231 | 0.357 | 0.722 | -0.065 | 0.799 | 0.997 |
| PCC | 0.541 | 0.020<sup>†</sup> | 0.286 | -0.441 | 0.067 | 0.623 |
| LIPL | 0.224 | 0.371 | 0.722 | -0.217 | 0.387 | 0.722 |
| RIPL | 0.158 | 0.533 | 0.785 | -0.019 | 0.942 | 0.997 |
| dACC | 0.169 | 0.502 | 0.785 | 0.235 | 0.349 | 0.722 |
| LAmy | 0.007 | 0.979 | 0.997 | 0.044 | 0.861 | 0.997 |
| RAmy | -0.334 | 0.176 | 0.703 | 0.384 | 0.115 | 0.646 |
| **fALFF** |  |  |  |  |  |  |
| mPFC | -0.014 | 0.957 | 0.997 | -0.001 | 0.997 | 0.997 |
| PCC | 0.344 | 0.163 | 0.703 | -0.560 | 0.016<sup>†</sup> | 0.286 |
| LIPL | 0.129 | 0.609 | 0.853 | -0.260 | 0.297 | 0.722 |
| RIPL | 0.090 | 0.722 | 0.963 | 0.015 | 0.952 | 0.997 |
| dACC | -0.172 | 0.495 | 0.785 | 0.233 | 0.353 | 0.722 |
| LAmy | 0.237 | 0.343 | 0.722 | -0.222 | 0.377 | 0.722 |
| RAmy | -0.165 | 0.513 | 0.785 | 0.385 | 0.115 | 0.646 |

Note: *adj\_p*, adjusted *p* value through FDR correction. \*, results survived $p < 0.05$ FDR correction. <sup>†</sup>, $p < 0.05$ without correction. The partial correlation between ALFF/fALFF and one variable of interest was conducted while controlling the effects of all the other variables non-investigated. mPFC, medial prefrontal cortex; PCC, posterior cingulate cortex; L/R IPL, left/right inferior parietal lobule; dACC, dorsal anterior cingulate cortex; L/R Amy, left/right amygdala.

For **Model II**, partial correlations between MIES-transgression/-betrayal or CAPS scores and average ALFF/fALFF in ROIs are reported in **Table 3**. ALFF in the left IPL was positively related with MIES-transgression (*R* = 0.776, *p* = 0.008 FDR corrected, **Fig. 2A**), negatively related with MIES-betrayal (*R* = −0.759, *p* = 0.008 FDR corrected, **Fig. 2B**), and has no relationship with CAPS scores (*R* = −0.337, *p* = 0.615 FDR corrected). Moreover, Williams’s t-tests showed that the ALFF correlation with MIES-transgression was significantly larger than the correlation with MIES-betrayal (*t* = 8.188, *p* < 0.001) and the correlation with CAPS (*t* = 7.852, *p* < 0.001). The ALFF correlation with MIES-betrayal was not significantly different from the correlation with CAPS (*t* = −2.090, *p* = 0. 976). No fALFF results survived FDR correction in Model II.

**Table 3.** Partial correlations between MIES-transgression/-betrayal and average ALFF/fALFF in ROIs.

| ROI | <u>MIES-transgression</u> |  |  | <u>MIES-betrayal</u> |  |  | <u>CAPS</u> |  |  |
| --- | --- | --- | --- | --- | --- | --- | --- | --- | --- |
|  | *R* | *p* | *adj p* | *R* | *p* | *adj p* | *R* | *p* | *adj p* |
| **ALFF** |  |  |  |  |  |  |  |  |  |
| mPFC | 0.314 | 0.220 | 0.615 | -0.103 | 0.695 | 0.912 | -0.060 | 0.818 | 0.954 |
| PCC | 0.337 | 0.186 | 0.615 | 0.468 | 0.058 | 0.334 | -0.459 | 0.064 | 0.334 |
| LIPL | 0.776 | 0.000<sup>†</sup> | 0.008<sup>*</sup> | -0.759 | 0.000<sup>†</sup> | 0.008<sup>*</sup> | -0.337 | 0.186 | 0.615 |
| RIPL | 0.210 | 0.419 | 0.710 | -0.061 | 0.815 | 0.954 | -0.015 | 0.954 | 0.983 |
| ACC | 0.315 | 0.218 | 0.615 | -0.201 | 0.440 | 0.710 | 0.252 | 0.329 | 0.710 |
| LAmy | -0.135 | 0.606 | 0.877 | 0.207 | 0.425 | 0.710 | 0.040 | 0.880 | 0.983 |
| RAmy | -0.492 | 0.045<sup>§</sup> | 0.314 | 0.219 | 0.399 | 0.710 | 0.404 | 0.108 | 0.505 |
| **fALFF** |  |  |  |  |  |  |  |  |  |
| mPFC | 0.165 | 0.526 | 0.789 | -0.261 | 0.313 | 0.710 | 0.007 | 0.980 | 0.983 |
| PCC | 0.206 | 0.428 | 0.710 | 0.265 | 0.305 | 0.710 | -0.565 | 0.018<sup>†</sup> | 0.180 |
| LIPL | 0.553 | 0.021<sup>†</sup> | 0.180 | -0.577 | 0.015<sup>†</sup> | 0.180 | -0.316 | 0.217 | 0.615 |
| RIPL | 0.093 | 0.723 | 0.915 | 0.006 | 0.983 | 0.983 | 0.016 | 0.950 | 0.983 |
| ACC | 0.006 | 0.982 | 0.983 | -0.281 | 0.274 | 0.710 | 0.245 | 0.344 | 0.710 |
| LAmy | 0.179 | 0.492 | 0.765 | 0.118 | 0.652 | 0.883 | -0.222 | 0.392 | 0.710 |
| RAmy | -0.120 | 0.647 | 0.883 | -0.087 | 0.740 | 0.915 | 0.386 | 0.127 | 0.531 |

Note: *adj\_p*, adjusted *p* value through FDR correction. <sup>\*</sup>, results survived *p* < 0.05 FDR correction. <sup>†</sup>, *p* < 0.05 without correction. The partial correlation between ALFF/fALFF and one variable of interest was conducted while controlling the effects of all the other variables non-investigated. mPFC, medial prefrontal cortex; PCC, posterior cingulate cortex; L/R IPL, left/right inferior parietal lobule; dACC, dorsal anterior cingulate cortex; L/R Amy, left/right amygdala.

**Figure 2.**
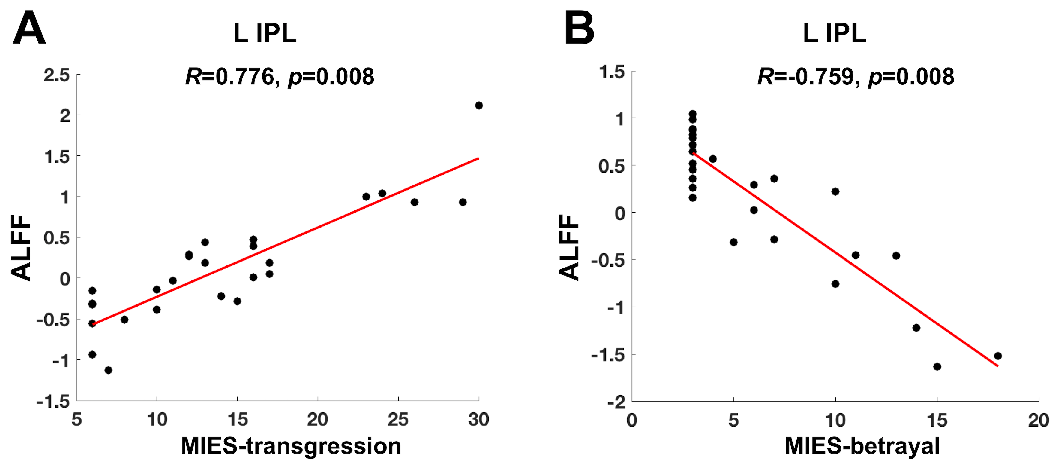
ALFF partial correlations. Larger ALFF In the ROI of left inferior parietal lobule (L IPL) was associated with (A) higher scores of moral injury transgression (MIES-transgression) (*R* = 0.776, *p* = 0.008 FDR corrected) and (B) lower scores of moral injury betrayal (MIES-betrayal) (*R* = −0.759, *p* = 0.008 FDR corrected). The mean ALFF values in the scatter plots are adjusted to regress out the effects of all the other variables non-investigated.

### ALFF and fALFF Whole-Brain Analyses Findings

For **Model I**, MIES-total was negatively related with ALFF in the right posterior insula (maximum effect at x/y/z/ = 38/−20/20, Z value = 4.57, cluster size = 132 voxels).

For **Model II**, as shown in **Table 4**, MIES-transgression was positively related with ALFF in the left IPL (**Fig. 3A**), and negatively related with ALFF in the right fusiform gyrus and right posterior insula. MIES-betrayal was positively related with ALFF in the left precuneus, and negatively related with ALFF in the left angular gyrus within left IPL (**Fig. 3B**) and right superior parietal lobule. Moreover, MIES-transgression was positively associated with fALFF in the left middle occipital gyrus and right angular gyrus in the right IPL. MIES-betrayal was positively related with fALFF in the right precuneus and right precentral gyrus.

**Table 4.** Whole-brain voxel-wise ALFF/fALFF correlations with MIES-transgression/-betrayal.

| Area | Size | Z | MNI |  |  |
| --- | --- | --- | --- | --- | --- |
|  |  |  | x | y | z |
| ALFF positively related with MIES-transgression |  |  |  |  |  |
| L Inferior Parietal Lobule (BA19/39) | 361 | 4.39 | -42 | -78 | 38 |
| ALFF negatively related with MIES-transgression |  |  |  |  |  |
| R Fusiform Gyrus (BA18/19) | 173 | 4.00 | 30 | -76 | -16 |
| R Posterior Insula (BA13) | 109 | 3.93 | 36 | -18 | 20 |
| ALFF positively related with MIES-betrayal |  |  |  |  |  |
| L Precuneus (BA7) | 112 | 4.22 | -14 | -54 | 54 |
| ALFF negatively related with MIES-betrayal |  |  |  |  |  |
| L Angular Gyrus (BA39) | 135 | 4.30 | -42 | -78 | 40 |
| R Superior Parietal Lobule (BA7) | 128 | 3.80 | 28 | -60 | 62 |
| fALFF positively related with MIES-transgression |  |  |  |  |  |
| L Middle Occipital Gyrus (BA19/39) | 117 | 4.44 | 44 | -76 | 36 |
| R Angular Gyrus (BA39/19) | 69 | 4.56 | -44 | -84 | 28 |
| fALFF negatively related with MIES-transgression |  |  |  |  |  |
| NS |  |  |  |  |  |
| fALFF positively related with MIES-betrayal |  |  |  |  |  |
| R Precuneus (BA7) | 93 | 4.54 | 16 | -62 | 48 |
| R Precentral Gyrus (BA6) | 70 | 4.42 | 48 | -2 | 28 |
| fALFF negatively related with MIES-betrayal |  |  |  |  |  |
| NS |  |  |  |  |  |

Note: all results were height-thresholded at $p < 0.001$ and survived $p < 0.05$ cluster-level FDR correction. BA, Brodmann's area; Size, number of voxels within the cluster; Z, z value; x/y/z, MNI coordinates. L, left, R, right.

**Figure 3.**
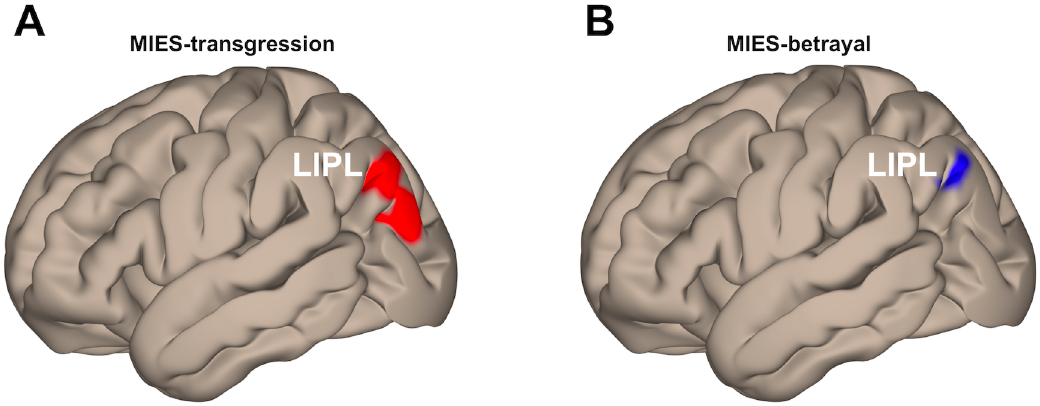
Whole-brain voxel-wise ALFF correlations. Larger ALFF in left inferior parietal lobule (LIPL) was associated with (A) higher scores of moral injury transgression (MIES-transgression, maximum effect at x/y/z/ = −42/−78/38) and (B) lower scores of moral injury betrayal (MIES-betrayal, maximum effect at x/y/z/ = −42/−78/40). Results were height-thresholded at *p* < 0.001 uncorrected and survived *p* < 0.05 cluster-extent level FDR correction.

### Seed-Based Whole-Brain Functional Connectivity Analyses Results

For **Model I**, MIES-total was positively correlated with functional connectivity between right amygdala seed and right thalamus (maximum effect at x/y/z/ = 14/−34/6, Z value = 4.73, cluster size = 214 voxels). No significant relationship was detected between CAPS and functional connectivity to the same seed-target pair.

Moreover, CAPS was positively correlated with functional connectivity between left IPL seed and bilateral precuneus (maximum effect at x/y/z/ = −10/−54/52, Z value = 4.21, cluster size = 461 voxels, **Fig. 4**). Further analyses showed that MIES-total was negatively correlated with functional connectivity to the same seed-target pair (maximum effect at x/y/z/ = −10/−52/52, Z value = 3.28, cluster size = 8 voxels, SVC).

**Figure 4.**
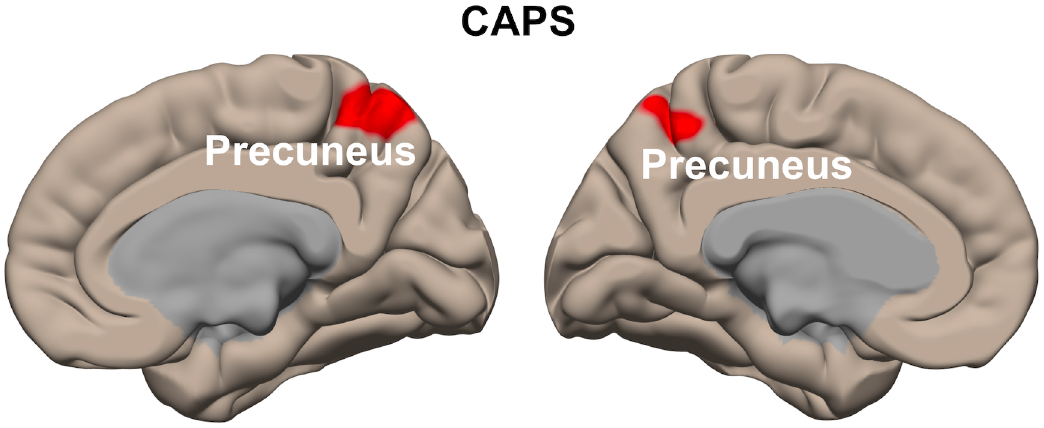
Whole-brain seed-based functional connectivity correlations. CAPS was positively correlated with functional connectivity between left IPL seed and left precuneus (maximum effect at x/y/z/ = −10/−54/52) as well as right precuneus (maximum effect at x/y/z/ = 6/−50/58). The MIES-total was negatively correlated with functional connectivity to the same seed-target pair.

For **Model II**, MIES-transgression was positively correlated with functional connectivity between left amygdala seed and left fusiform gyrus (maximum effect at x/y/z/ = −34/−42/−30, Z value = 4.56, cluster size = 250 voxels). Further analyses showed that both MIES-betrayal (maximum effect at x/y/z/ = −34/−40/−32, Z value = 4.11, cluster size = 20 voxels, SVC) and CAPS (maximum effect at x/y/z/ = −42/−54/−26, Z value = 3.92, cluster size = 40 voxels, SVC) were negatively correlated with functional connectivity for the same seed-target pair.

Moreover, CAPS was positively correlated with functional connectivity between left IPL seed and left precuneus (maximum effect at x/y/z/ = −10/−54/52, Z value = 4.08, cluster size = 212 voxels) and right precuneus (maximum effect at x/y/z/ = 6/−50/58, Z value = 3.84, cluster size = 210 voxels). Neither MIES-transgression nor MIES-betrayal was found to correlate with functional connectivity between left IPL seed and precuneus.

## Discussion

The present study examines the neural correlates of moral injury and PTSD, as well as the neural correlates of two moral injury sub-scales, transgression and betrayal, in combat veterans. We found that ALFF in the left IPL was positively related with MIES-transgression, negatively associated with MIES-betrayal, and had no relationship with PTSD. Moreover, functional connectivity between the left IPL and bilateral precuneus was positively related with PTSD symptoms and negatively related with total scores of moral injury. These results mark the left IPL as a locus of dissociable neural correlates between moral injury and PTSD, as well as a location of distinct brain responses to moral injury from transgressive acts and moral injury from betrayal.

Both ROI and whole-brain analyses on ALFF uncovered the neural correlates of moral injury sub-scales, i.e. MIES-transgression and MIES-betrayal, in the left IPL. The IPL serves as a major hub for integrating multi-sensory information inputs for comprehension and manipulation [43]. It is also an important component of both the DMN [13] that plays a role in internally directed or self-generated thoughts, and the ToM regions, which infer the mental states of others [44]. This area robustly activates during tasks of responding to moral dilemmas, violations of moral principles, and making moral decisions [45]. Accordingly, there is reduced glucose metabolism [46] and decreased regional cerebral blood flow [47] in IPL of criminals convicted of impulsive violence and murders. These previous studies highlight the role of IPL in social cognition and moral processing, consistent with our findings of the significant relationships between moral injury and resting-state brain responses. By contrast, we did not find any significant relationship between ALFF (or fALFF) in IPL and CAPS scores. This negative finding is inconsistent with a recent meta-analysis reporting that spontaneous brain activity in the left IPL is positively correlated with PTSD symptom severity [32]. A potential explanation is that the aforementioned meta-analysis on fear-based PTSD did not clarify the potential biases incurred by moral injury, shame, guilt or disrupted social cognition that are often accompanied with PTSD symptoms [1; 7]. The exact role of resting-state spontaneous fluctuations in the left IPL still needs further investigation in PTSD.

It is also interesting that ALFF in the left IPL was positively correlated with transgression scores, and negatively correlated with betrayal scores. This finding suggests distinct neural underpinnings in left IPL between perceived transgression and betrayal, consistent with a previous behavioral study [8] that dissociated the two latent factors in veterans suffering from moral injury. However, it is hard to determine the exact alteration in neural processing in IPL solely based on ALFF, given the complicated relationship between resting-state brain responses and task-related brain activations. Firstly, stronger ALFF may be related to larger task-related activation in some areas, but smaller activation in other areas [22]. Secondly, larger ALFF does not necessarily represent more efficient processing but a compensatory effect for deficits in patients with specific disorders [48]. Thirdly, beyond social cognition, the IPL is associated with semantic processing, number processing, memory retrieval, spatial attention and reasoning [49]. Resting-state data cannot differentiate multiple functions in the same area and therefore the exact role of the left IPL in moral injury needs to be clarified with task-based neuroimaging studies. The IPL is more active when engaged in tasks evaluating moral dilemmas [50], but less active when posed with moral conflicts as compared to analogous non-moral scenarios [51]. A recent study on moral transgression in healthy participants by Crockett et al. [52] developed a task paradigm in which non-clinical participants made decisions whether to accrue monetary benefits by inflicting pains on others. Future neuroimaging studies employing this paradigm or similar tasks may help to directly investigate the neural responses to moral dilemmas and social decisions in people suffering from moral injury.

Besides the ALFF findings, we found that functional connectivity between left IPL and bilateral precuneus was positively related with PTSD symptoms and negatively associated with moral injury total scores, providing further support of the neural dissociations between moral injury and PTSD. The IPL-precuneus functional connections have been reported in previous rs-fMRI studies [53]. Task-based studies have also documented the co-activations of IPL and precuneus in attention, self-perception, introspection and memory, and social cognition [54]. It is possible that moral injury and PTSD are different in a few of these cognitive functions. However, this idea needs to be tested in task-based studies.

Our findings may contribute to designing clinical intervention of moral injury and PTSD. Resting-state activity has been reported to predict behavioral performance [22]. Therefore, spontaneous neural activity in left IPL and functional connectivity between left IPL and precuneus may be utilized to complement the assessments of moral injury/PTSD and monitoring the response to clinical intervention [55]. Moreover, increasing numbers of studies have documented training-induced plasticity in bilateral IPL [56]. Learning new skills on spatial coordination, verbal memory, and emotional regulation practices were found to increase grey matter volume in IPL [57]. Given the multiple roles of IPL in not only social cognition, but also spatial and verbal processing [49], one might hypothesize that training to enhance spatial and/or verbal abilities may modulate brain structures including IPL, which may have collateral benefits for treating moral injury/PTSD. This idea would certainly require testing in future studies.

There are a few limitations in the present study. Firstly, the correlation and regression models utilized here may overlook the non-linear relationships between clinical measures and brain responses. Future studies aimed at dissociating the neural correlates of moral injury and PTSD may consider comparing four groups of participants: (1) PTSD without moral injury, (2) trauma-exposed controls without PTSD nor moral injury, (3) high scores in moral injury without PTSD, and (4) low scores in moral injury without PTSD. Thus, the contrast between group 1 and 2 will unveil the neural correlates of PTSD, while the comparison between group 3 and 4 will uncover the neural underpinnings of moral injury. A second potential limitation is that the MIES-transgression sub-score includes a mix of exposures (questions 1, 3 and 5 in MIES) and symptoms (questions 2, 4, and 6 in MIES), whereas the MIES-betrayal sub-score pertains only to symptoms. Despite the results of a factor analysis by Nash and colleagues [8], which did not reveal separate components for transgression-related exposures and transgression-related symptoms, we nevertheless reasoned that these two constructs might have distinct neural correlates. Our analyses (see Supplementary Materials) demonstrated that, ALFF in the left IPL was positively correlated with transgression-related symptoms as well as exposures, supporting the validity of the main findings. Future studies on moral injury with an improved classification will help to delineate the neurobiological subtypes of moral injury.

In conclusion, we found that PTSD and moral injury subscales, i.e. transgression and betrayal, are dissociated by the ALFF in left IPL. Moreover, PTSD and moral injury total scores are differentiated by the functional connectivity between left IPL and precuneus. Our findings significantly enhance our understanding of the neural correlates of moral injury vis-à-vis PTSD, and shed light on neural targets for potential clinical interventions. Knowledge of relevant targets could help predict, guide selection, or monitor treatment response of psychotherapy, pharmacotherapy, or brain stimulation, which may be optimally suited for individual patients. To the best of our knowledge, this is the first publication uncovering the neural correlates of moral injury, and the first study that documents the neural differences between moral injury and PTSD. Moral injury offers a complementary behavioral model that extends prevailing fear and threat models of PTSD [58]. Expanding our investigation into the neuroscience of moral processing may open new avenues of research that enrich our understanding of PTSD beyond the existing fear-based models.

## Acknowledgements

This research was supported by the U.S. Department of Veterans Affairs (VA) Mid-Atlantic Mental Illness Research, Education, and Clinical Center (MIRECC) core funds and financial support by the Duke Health Scholars Award to Dr. Morey. Dr. Sun was supported by the NIH (5R01-NS086885) and Mid-Atlantic MIRECC pilot research funds. We thank Mira Brancu PhD and Kevin LaBar PhD for their helpful comments and suggestions in the early versions of the manuscript.

